# A global co-expression network approach for connecting genes to specialized metabolic pathways in plants

**DOI:** 10.1101/093914

**Authors:** Jennifer H. Wisecaver, Alexander T. Borowsky, Vered Tzin, Georg Jander, Daniel J. Kliebenstein, Antonis Rokas

## Abstract

Plants produce a tremendous diversity of specialized metabolites (SMs) to interact with and manage their environment. A major challenge hindering efforts to tap this seemingly boundless source of pharmacopeia is the identification of SM pathways and their constituent genes. Given the well-established observation that the genes comprising a SM pathway are co-regulated in response to specific environmental conditions, we hypothesized that genes from a given SM pathway would form tight associations (modules) with each other in gene co-expression networks, facilitating their identification. To evaluate this hypothesis, we used 10 global co-expression datasets—each a meta-analysis of hundreds to thousands of expression experiments—across eight plant model organisms to identify hundreds of modules of co-expressed genes for each species. In support of our hypothesis, 15.3-52.6% of modules contained two or more known SM biosynthetic genes (e.g., cytochrome P450s, terpene synthases, and chalcone synthases), and module genes were enriched in SM functions (e.g., glucoside and flavonoid biosynthesis). Moreover, modules recovered many experimentally validated SM pathways in these plants, including all six known to form biosynthetic gene clusters (BGCs). In contrast, genes predicted based on physical proximity on a chromosome to form plant BGCs were no more co-expressed than the null distribution for neighboring genes. These results not only suggest that most predicted plant BGCs do not represent genuine SM pathways but also argue that BGCs are unlikely to be a hallmark of plant specialized metabolism. We submit that global gene co-expression is a rich, but largely untapped, data source for discovering the genetic basis and architecture of plant natural products, which can be applied even without knowledge of the genome sequence.

## Introduction

Plants, being sessile and therefore at the mercy of their surroundings, harbor many adaptations that facilitate their interaction with and management of their environment. One such adaptation is the ability to produce a vast array of specialized metabolites (SMs), bioactive compounds that are not essential for growth and reproduction but rather have important ecological roles to combat pathogens, herbivores, and competitors; attract pollinators and seed dispersers; and resist abiotic stress including fluctuations in temperature, salinity, and water availability^1^. Humans exploit the SM diversity of plants for medicines and other natural products; to this end, thousands of plant-derived SMs have been isolated and biochemically characterized^2^. Yet the genes responsible for the production and regulation of most SMs across the kingdom Plantae are unknown, which ultimately limits their potential utility in agricultural, pharmaceutical, and biotechnological applications^3,4^.

Given their biomedical and agricultural relevance, it is perhaps surprising that the constituent genes and pathways involved in biosynthesis of most plant SMs are unknown^5^. There are two explanations for why this is so; first, SM pathways are highly variable in the number and functions of genes they contain^1,6^. Second, consistent with their involvement in the production of ecologically specialized bioactive molecules, SM genes exhibit narrow taxonomic distributions, are fast evolving both in terms of sequence divergence and rate of gene family diversification, and display extensive functional divergence^7–9^. The consequence of this lack of evolutionary and functional conservation is that traditional sequence homology metrics for inferring gene function^10^ are weak predictors of SM pathway composition and function.

Network biology offers a promising alternative for identifying SM pathways and their constituent genes. Because SM pathways exist at the interface of organisms and their environments, the genes within an SM pathway share a common regulatory network that tightly controls the “where” (e.g., in what tissues) and “when” (e.g., in response to which ecological conditions) of SM production^1,11,12^. Therefore, gene co-expression data, as a proxy for co-regulation, have been particularly effective in identifying the constituent genes that make up many SM pathways^13–21^. Further, given the availability of data from hundreds to thousands of individual gene expression experiments, integrative global co-expression networks have the power to predict SM pathways and genes in a high-throughput fashion^22–24^. However, as measuring gene co-expression on a large scale was, until recently, a costly and labor-intensive undertaking, the hundreds (or more) of global gene expression studies in diverse conditions required for global co-expression network analyses currently exist for only a small minority of plant species^25–27^.

Another attribute that is characteristic of SM pathways found in bacteria and fungi is that they can physically co-locate in the genome, forming biosynthetic gene clusters (BGCs)^28^. As expected of SM pathways, genes within these microbial BGCs are co-regulated and display strong signatures of co-expression, a pattern that holds true for functionally characterized as well as for putative BGCs in these genomes^29–33^. As the proximity of genes on chromosomes is far easier to measure than their co-expression across multiple experimental conditions, bioinformatic algorithms strongly rely this “clustering” of genes to predict SM pathways in microbial genomes^34–36^. Thus, thousands of microbial BGCs have been predicted and hundreds validated (i.e., connected to known products), suggesting that gene proximity is informative for SM pathway identification, at least in these organisms^37^. Nevertheless, the number of SM pathways in bacteria and fungi that do not (or only partially) form BGCs is unknown^38–40^.

In plants, most characterized SM pathways (e.g., glucosinolate biosynthesis) are not clustered, and their genes are distributed across the genome^41^. More recently, however, nearly two dozen BGCs responsible for the production of SM defensive compounds have been identified and functionally characterized from 15 plant species^42^, raising the possibility that gene proximity could also be used for predicting plant SM pathways^43^. To this end, computational searches based on gene clustering similar to those developed for fungal and bacterial genomes postulate the existence of dozens to hundreds more BGCs across a wide variety of plant genomes^8,44,45^. However, the vast majority of these putative plant BGCs has not been functionally validated, and the fraction of plant SM pathways that form BGCs is unclear.

We hypothesized that plant SM pathways are co-expressed, independently of being organized into BGCs, in line with their ecological roles that require strong temporal and spatial co-regulation^1,11,12^. To test our hypothesis, we developed a gene co-expression network-based approach for plant SM pathway discovery (Figure S1) using data from 10 meta-analyses of global co-expression that collectively contain 21,876 microarray or RNA-Seq experiments across eight plant species. Doing so, we identified dozens to hundreds of modules of co-expressed genes containing SM biosynthetic genes (e.g., cytochrome P450s, terpene synthases, and chalcone synthases) in each species, including many experimentally validated SM pathways and all validated BGCs in these species. In contrast, genes predicted to be in BGCs based on their physical proximity did not exhibit significantly different co-expression patterns than their non-clustered neighbors. Our results cast doubt on the general utility of approaches for SM pathway identification based on gene proximity in the absence of functional data and suggest that global gene co-expression data, when in abundance, are very powerful in the high-throughput identification of plant SM pathways.

## Results

### Network analysis identifies small, overlapping modules of co-expressed genes in global co-expression networks

Given that SM pathway genes are often co-regulated in response to specific environmental conditions, we hypothesized that genes from a given SM pathway would form tight associations (modules) with each other in gene co-expression networks. To identify modules of co-expressed SM genes, we accessed three microarray-and seven RNAseq-based co-expression datasets from ATTED-II^25^ and ALCOdb^46^ for eight Viridiplantae species (*Arabidopsis thaliana, Brassica rapa, Chlamydomonas reinhardtii, Glycine max, Oryza sativa* Japonica group, *Populus trichocarpa, Solanum lycopersicum,* and *Zea mays;* Table S1). Each dataset consisted of a meta-analysis of hundreds to thousands of experiments measuring global patterns of gene expression in a wide variety of tissues, environmental conditions, and developmental stages. The number of experiments varied in each dataset, from 172 in the *C. reinhardtii* RNAseq dataset^46^ to 15,275 in the *A. thaliana* microarray-based set^25^. Pairwise measurements of gene coexpression were specified as Mutual Ranks^47^ (MRs; calculated as the geometric mean of the rank of the Pearson’s correlation coefficient (PCC) of gene A to gene B and of the PCC rank of gene B to gene A). For each dataset, we constructed five MR-based networks, each using a different co-expression threshold for assigning edge weights (connections) between nodes (genes) in the network. Networks were ordered based on size (i.e., number of nodes and edges), such that N1 and N5 indicated the smallest and largest networks, respectively.

To discover co-expressed gene modules in the eight model plants, we employed the graph-clustering method ClusterONE^48^, which allowed genes to belong to multiple modules. This attribute is biologically realistic; many plant metabolic pathways are non-linear, containing multiple branch points and alternative end products (e.g., terpenoid biosynthesis pathways^49,50^). Averaging across all 10 co-expression datasets, the number of genes assigned to modules ranged from 3,251 (13.4% of protein-coding genes) in the N1 networks to 4,320 (18.2%) in N5 networks (Table S2). The average number of modules per network decreased with increasing network size, from 573 modules in the N1 networks to 39 in the N5 networks (Table S2). Conversely, the average module size (i.e., number of genes within a module) increased with increasing network size (e.g., 7 genes per module in N1 networks, 41 genes per module in N3 networks, and 167 genes per module in N5 networks). Given our goal to recover distinct SM pathways as modules, we focused the remaining analyses on the smaller networks (N1-N3) with average module sizes (< 50 genes) consistent with the number of genes typically present in SM pathways.

### Co-expressed gene modules recover known SM pathways and predict hundreds of new SM gene associations

To evaluate the correspondence between module genes and genes present in known metabolic pathways, we focused on the 798 genes in 362 *A. thaliana* MetaCyc^51^ pathways with an experimentally validated metabolic function (Table S3). Module genes were significantly enriched in many SM-related metabolic functions. Of the 12 higher-order metabolic classes investigated, only the SECONDARY METABOLITES and CELL STRUCTURES biosynthesis classes were significantly enriched in module genes (*P* < 0.0005, hypergeometric tests) (Figure 1a). This pattern held true across all networks and datasets investigated (Figure S2 and Table S4). Enrichment of the CELL STRUCTURES biosynthesis class was driven by genes involved in the SECONDARY CELL WALL (specifically LIGNIN) biosynthesis subclasses (P < 0. 0005, hypergeometric tests). Enriched subclasses within the SECONDARY METABOLITES class included those for NITROGEN-CONTAINING SECONDARY COMPOUNDS and FLAVONOID biosynthesis (P < 0.005, hypergeometric tests), which contain pathways for glucosinolate and anthocyanin production, respectively. MetaCyc SM pathways that were well recovered as coexpressed modules included those for aliphatic and indolic glucosinolate, camalexin, flavonol, flavonoid, phenylpropanoid, spermidine, and thalianol biosynthesis (Table S5).

**Figure 1.**
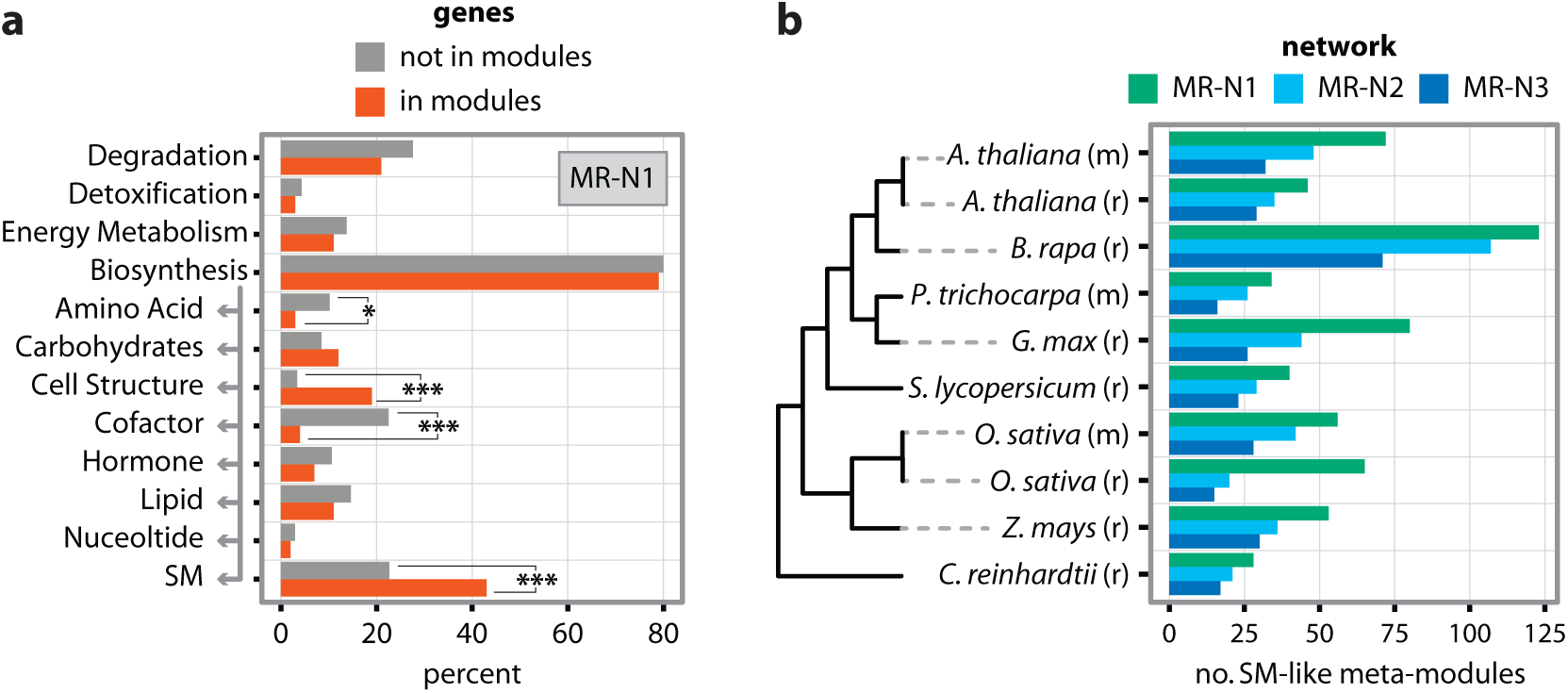
Global co-expression network analysis of eight plant genomes identify co-expressed modules of specialized metabolic genes. a) MetaCyc pathway enrichment analysis of experimentally characterized *A. thaliania* genes assigned to modules (orange bars) relative to those that do not form modules (grey bars) in *A. thaliania* microarray-based network MR-N1. Grey arrow indicates that the bottom eight pathway categories are children of ‘Biosynthesis’ in the MetaCyc hierarchy. Asterisks denote significant enrichment or depletion of MetaCyc categories in module genes; \**P* ≤ 0.05, \*\*\**P* ≤ 0.0005 (Benjamini & Hochberg adjusted *P*-values, hypergeometric test). See Figure S2 for enrichment tests in other *A. thaliania* networks. b) Count of SM-like meta-modules identified in 10 microarray (m) and RNAseq (r) co-expression datasets from eight Viridiplantae. SM-like modules were collapsed into meta-modules of non-overlapping gene sets. Networks were constructed using three different rates of exponential decay for converting MR scores to edge weights, where MR-N1 corresponds to smallest network with the steepest rate of decay and therefore the fewest edges; conversely, MR-N3 is the largest network with the shallowest rate of decay and the most edges.

The AMINO ACIDS, CARBOHYDRATES, and COFACTORS/PROSTHETIC GROUPS/ELECTRON CARRIERS biosynthesis classes were significantly depleted in module genes in some, but not all, networks and datasets (P < 0.05, hypergeometric test) (Figure 1a and Figure S2). None of the other metabolic classes displayed any significant variation between module and non-module genes (Figure S2 and Table S4).

To estimate the number of modules that may correspond to SM pathways, we focused on those that contained two or more non-homologous genes with a significant match to a curated list of PFAM domains that are found commonly in genes from SM pathways (Table S6); as some of these “SM-like” modules share genes, we collapsed them into non-intersecting “meta-modules”. Dozens of SM-like meta-modules were identified in each species, with the green alga, *C. reinhardtii,* containing the fewest SM-like meta-modules (27 in N1 networks, 17 in N3 networks), and the field mustard, *B. rapa,* containing the most (120 in N1 networks, 71 in N3 networks) (Figure 1b and Table S2).

### Recovery of the aliphatic glucosinolate biosynthesis pathways in *Arabidopsis* and *Brassica* from global co-expression data

To illustrate the utility and power of our approach for identifying entire SM pathways, we next focused on examining the correspondence between genes involved in the methionine-derived aliphatic glucosinolate (metGSL) biosynthesis pathway and genes that comprise co-expression modules identified by our analyses (Table S7). In *A. thaliana,* the species with the majority of functional data^52^, co-expression modules recover genes for every biochemical step in this pathway, from methionine chain elongation to side-chain modification of the glucosinolate chemical backbone, as well as a pathway-specific transporter and three transcription factors (Figure 2a). For example, in the smallest network N1, 14 / 34 enzymatic genes in the metGSL pathway are recovered in a single 17-gene module; only 3 / 17 genes in this module have not been functionally characterized as involved in metGSL biosynthesis (Figure 2b). Maximum recovery of the metGSL pathway increased to 56.3% and 71.9% in the 22-gene and 43-gene modules recovered from networks N2 and N3, respectively (Figure S3 and Table S8). Although the numbers of genes not known to be involved in metGSL biosynthesis also increased in these modules, several of the genes that are co-expressed with members of the metGSL pathway perform associated biochemical processes (Figure 2a). For example, the two adenosine-5’- phosphosulfate kinase genes, *APK1* and *APK2,* are responsible for activating inorganic sulfate for use in the metGSL pathway and polymorphisms in these genes alter glucosinolate accumulation^53^. Similarly, the cytochrome P450 genes, *CYP79B2* and *CYP79B3,* and the glutathione S-transferase gene, *GSTF9,* are involved in the parallel pathway for biosynthesis of glucosinolates from tryptophan instead of methionine (MetaCyc PWY-601)^52^.

**Figure 2.**
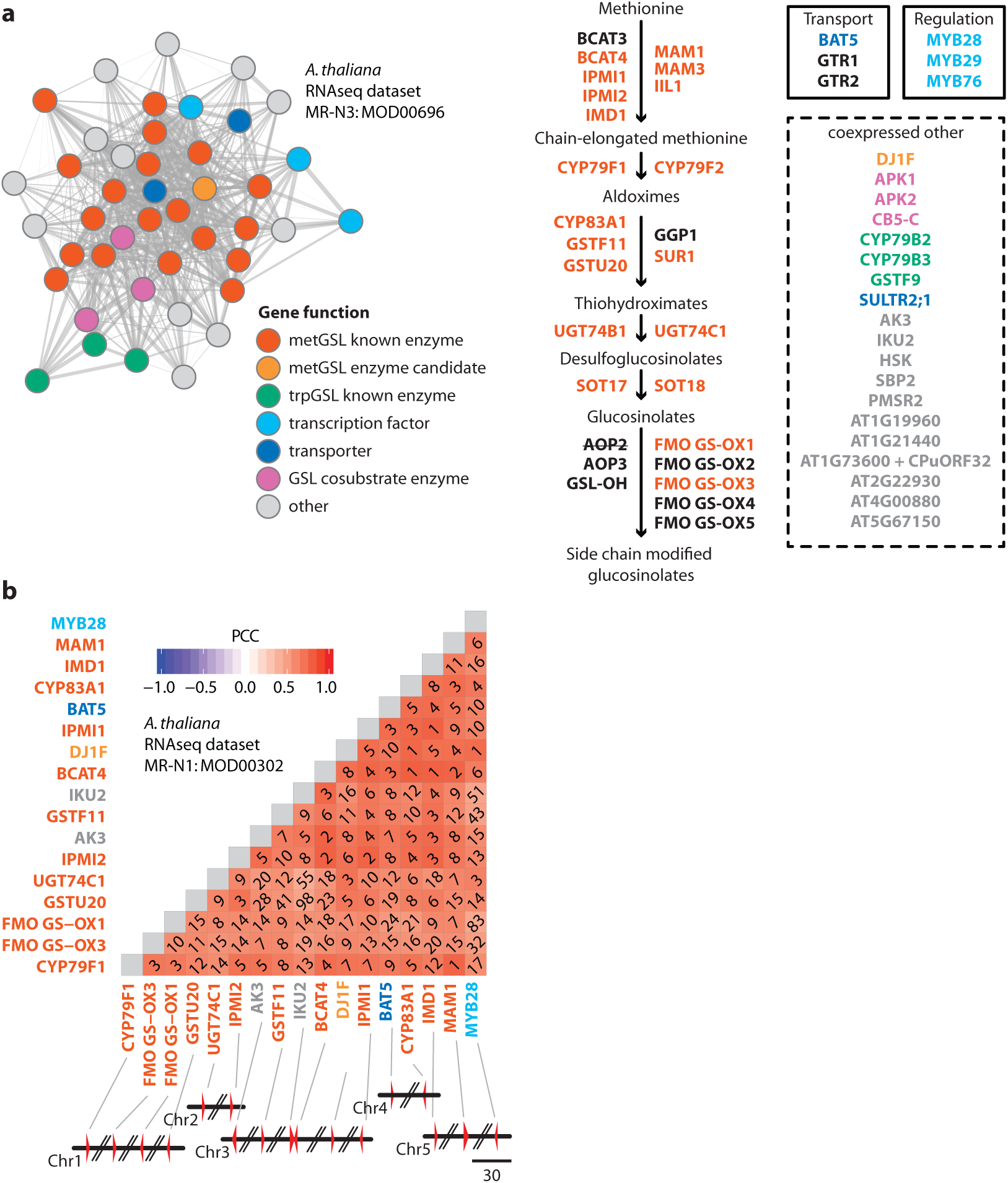
Co-expression modules efficiently recover the majority of genes for metGSL biosynthesis in *A. thaliana.* a) Network map of an example co-expression module involved in metGSL biosynthesis. Nodes in the map represent genes and edges connecting two genes represent the weight (transformed MR score) for the association. The diagram of the metGSL biosynthesis pathway is depicted, right. Other co-expressed genes not known to be involved in metGSL biosynthesis are shown in the dashed box. Nodes and gene names are colored according to their known or putative function. MetGSL genes not recovered in modules are colored black. Genes not present in the co-expression dataset are lined out. b) Heatmap depicting the correlation of co-expression of a second example co-expression module involved in metGSL biosynthesis. Diagonal numbers within the heatmap indicate MR score. Gene names are colored as in part a. Module genes are depicted as red triangles in the accompanying chromosome segments (parallel lines indicate the genes are not physically co-located; scale bar is in kilobase pairs). Note: data from the RNAseq-based networks are shown as two metGSL genes *(SOT17* and *SOT18)* are not present in the microarray dataset. Microarray-based networks performed similarly (Table S8).

Notably, some genes implicated in metGSL biosynthesis were never recovered in co-expressed modules, including *GGP1,* which encodes a class I glutamine amidotransferase-like protein. Microarray-based co-expression data weakly associate *GGP1* with metGSL biosynthesis in *A. thaliana,* and *GGP1* has been shown to increase glucosinolate production when heterologously expressed in *Nicotiana benthamiana^54^*. However, our metGSL-containing modules across all RNAseq-based networks showed that a different class I glutamine amidotransferase-like gene, *DJ1F*, is more highly co-expressed with metGSL biosynthetic genes (Figure 2). Importantly, *DJ1F* is not represented on the *A. thaliana* Affymetrix GeneChip, explaining why *GGP1* and not this gene was identified as the most correlated one in earlier analyses. However, the postulated role of both *DJ1F* and *GGP1* in metGSL biosynthesis remains to be confirmed *in planta.*

The remaining genes in the metGSL pathway that were never recovered in co-expressed modules all encode secondary enzymes responsible for terminal modifications to the backbone glucosinolate product^41^. One of these, *AOP2,* encoding a 2-oxoglutarate-dependent dioxygenase, has been pseudogenized in the *A. thaliana* (ecotype: Columbia) reference genome^55^. The high level of natural variation present in these terminal metabolic branches is responsible for the diverse glucosinolates present in different ecotypes^56,57^ but likely also makes it more challenging to connect them to the rest of the metGSL pathway using global co-expression data (Figure S4).

Brassicas also produce aliphatic glucosinolates, but a whole genome triplication event subsequent to their divergence from *A. thaliana*^58^ has complicated identification of functional metGSL genes in these species. To gain insight into the metGSL pathway in *B. rapa,* we cross-referenced our co-expression modules with 59 candidate metGSL genes identified based on orthology to *A. thaliana* metGSL genes^59^. As in *A. thaliana,* modules recovered every biochemical step of the *B. rapa* metGSL pathway as well as pathway-specific transporters and transcription factors (Figure 3, Table S7, and Table S8). Also as in *A. thaliana, DJ1F* rather than *GGP1* is co-expressed with other metGSL genes, providing further evidence that the DJ1F enzyme may be the more likely candidate for the γ-glutamyl peptidase activity in glucosinolate biosynthesis^52^. Furthermore, as several enzymes are encoded by multiple gene copies in *B. rapa*, we harnessed the power of our module analysis to identify which of these copies was co-expressed with other metGSL genes and therefore most likely to be functionally involved in the pathway. For example, out of the six *MAM* gene copies in *B. rapa,* only *Bra029355* and *Bra013007*were recovered in metGSL modules (Figure 3 and Figure S5). Module data also suggest that the glutathione S-transferase class tau (GSTU) activity is one step of the core pathway that may differ between the two species. Specifically, in *A. thaliana*, *GSTU20* is thought to encode this reaction, and this gene was recovered in metGSL modules in our analysis (Figure 2a). However, this association was not recovered in *B. rapa.* Instead, three paralogous GSTUs *(Bra003647, Bra026679,* and *Bra026680),* corresponding to the *A. thaliana GSTU23* and *GSTU25* genes, respectively, formed modules with metGSL genes, making these genes good candidates for investigation of GSTU activity in *B. rapa* (Figure 3 and Figure S6).

**Figure 3.**
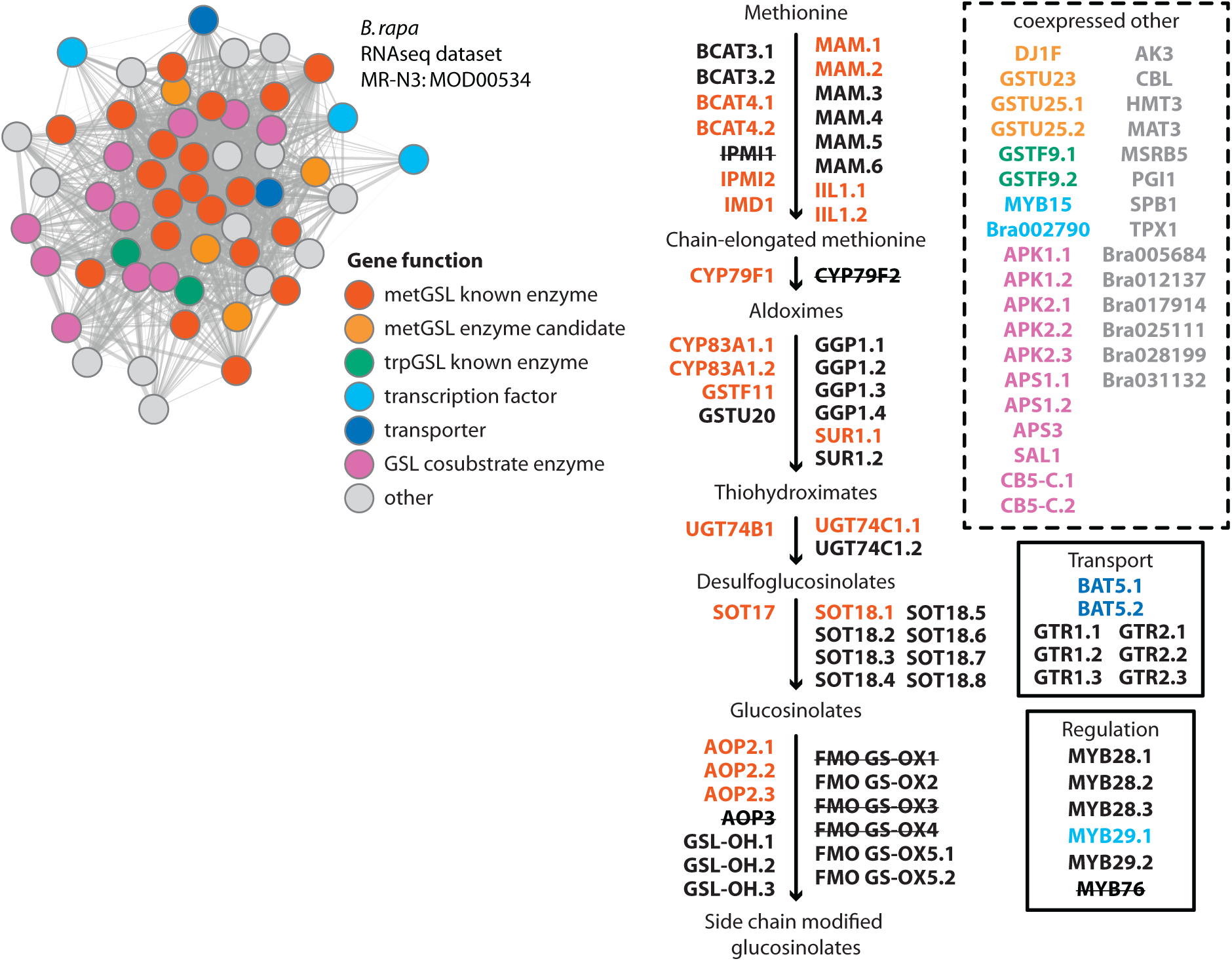
Co-expression modules predict functional metGSL biosynthesis genes in *B. rapa.* Network map of an example co-expression module involved in metGSL biosynthesis. Nodes in the map represent genes and edges connecting two genes represent the weight (transformed MR score) for the association. The diagram of the metGSL biosynthesis pathway is depicted, right, with all predicted orthologs to known metGSL genes in *A. thaliana* as listed on brassicadb.org. Other co-expressed genes not known to be involved in metGSL biosynthesis are shown in the dashed box. Nodes and gene names are colored according to their known or putative function. MetGSL orthologs not recovered in modules are colored black. *A. thaliana* metGSL genes with no known ortholog in *B. rapa* are lined out. See Table S7 for associated NCBI and Ensembl gene IDs.

### Modules recover functionally characterized BGCs and identify associated, unclustered genes

We next investigated whether our approach also recovered BGCs by examining whether our co-expression modules recovered the six functionally characterized BGCs in these eight plant genomes (Table S9). All six BGCs were recovered in our module analysis (Table S8). Specifically, co-expression modules recovered all genes comprising the BGCs involved in the production of the triterpenoids marneral^60^ (3 / 3 genes; Figure 4a) and thaliaol^61^ (4 / 4 genes; Figure 4b) in *A. thaliana* and the diterpenoid momilactone^62^ (5 / 5 genes; Figure 4c) in *O. sativa.* Modules recovered 7 / 9 genes in the phytocassane^63^ diterpene cluster in *O. sativa* (rice); the *OsKSL5* and *CYP71Z6* genes forming a terpene synthase-cytochrome p450 pair of genes were strongly co-expressed with each other but not with the rest of the pathway (Figure 4d). The two triterpenoid BGCs in *A. thaliana* were typically combined into the same co-expression module; the same pattern was observed for the two diterpenoid BGCs in *O. sativa* (Figure S7 and Figure S8). Genes within these BGCs were also strongly co-expressed with additional genes located outside the BGC boundaries, including one putative transcription factor and several putative transporters (Figure S7 and Figure S8).

**Figure 4.**
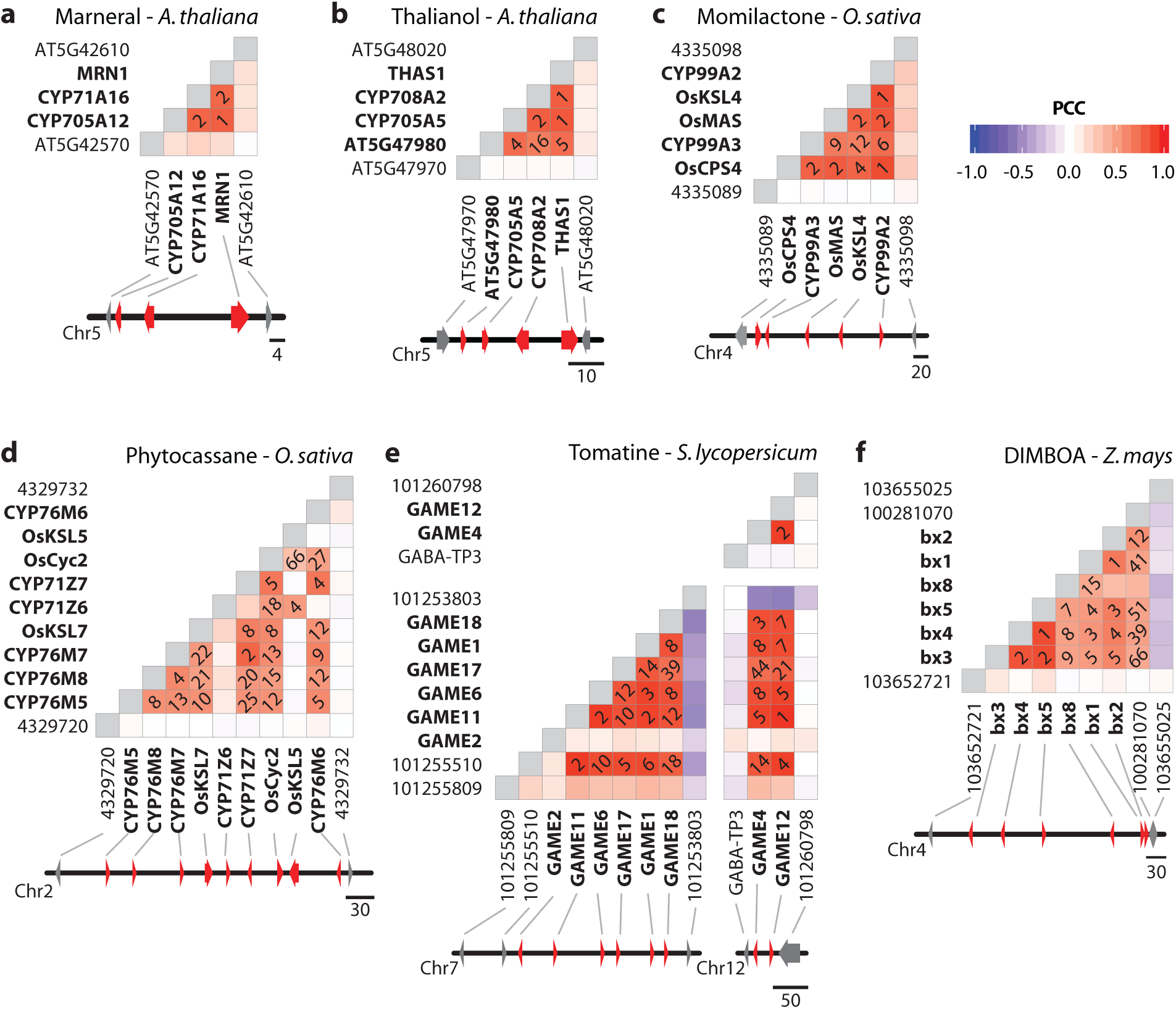
Co-expression pattern of six functionally characterized BGCs in plants. Heatmaps depict the correlation of co-expression of six BGCs for the production of a. marneral, b. thalianol, c. momilactone, d. phytocassane, e. tomatine, and f. DIMBOA. Diagonal numbers indicate MR scores; squares are blank if MR ≥ 100. BGC genes are bolded in the heatmap and colored red in the accompanying chromosome segments. Scale bars are in kilobase pairs.

Seven of eight genes in the partially clustered pathway for production of the steroidal alkaloid α-tomatine in *S. lycopersicum^64^* (tomato) were recovered by our coexpression analysis (Figure 4e). Only the glucosyltransferase gene, *GAME2,* encoding the last enzymatic reaction in the proposed α-tomatine pathway, showed a conspicuously different expression profile, consistent with previous reports^64,65^. Several glucosyltransferase genes paralogous to *GAME2* were strongly co-expressed with the rest of the genes in this pathway (Figure S9), but whether or not these genes participate in α-tomatine biosynthesis is yet to be determined. Additional genes strongly co-expressed with the rest of the α-tomatine pathway include, among others, one putative transcription factor, several possible metabolite transporters, and a cellulose synthase-like gene located adjacent to the BGC (Figure 4e and Figure S9).

Lastly, five of the six genes in the benzoxazinoid 2,4-dihydroxy-7-methoxy-1,4-benzoxazin-3-one (DIMBOA)^66^ cluster in *Z. mays* formed co-expression modules in our analysis (Figure 4f). Specifically, the first five genes in the DIMBOA pathway *(Bx1-Bx5),* responsible for the biosynthesis of the precursor 2,4-dihydroxy-1, 4-benzoxazin-3-one (DIBOA), formed modules with each other but not with the final gene in the BGC, *Bx8* (Figure S10).

Similar to the modifying genes of the metGSL pathway in *A. thaliana,* terminal *Bx* genes appear to have unique gene expression signatures distinct from the core pathway. For example, DIBOA is modified to DIMBOA by the action of two additional unclustered genes (*Bx6* and *Bx7*)^67^, neither of which was assigned to modules with core genes or each other. Toxic DIBOA/DIMBOA is transformed into stable glucoside, DIBOA-Glc/DIMBOA-Glc, by glucosyltransferases *(Bx8* and *Bx9),* which were likewise not assigned to modules in our analysis. However, a gene adjacent to the DIMBOA BGC, encoding an uncharacterized glucosyltransferase (GT; *GRMZM2G085854)* with 27% amino acid identity to *Bx8*, does belong to the same module as the core *Bx* genes in network N3 (Figure S10), but the MR scores of this gene to core *Bx* genes are noticeably weaker than those between the core *Bx* genes (Figure 4f). Additional *Bx* genes *(Bx10-Bx14),* which are not part of the BGC and are responsible for the biosynthesis of modified benzoxazinoid compounds (e.g., HDMBOA-Glc, DIM_2_BOA-Glc)^68,69^, were also not assigned to modules in our analysis (Figure S10); this pattern is similar to that observed with the terminal reactions of the metGSL biosynthesis pathway.

*Bx1* is thought to represent the first committed step in benzoxazinoid biosynthesis, encoding an indole-3-glycerolphosphate lyase (IGL) that converts indole-3-glycerolphosphate to indole. However, in our module analysis, an additional gene co-expressed with the core *Bx* genes is an indole-3-glycerolphosphate synthase (IGPS; *GRMZM2G106950),* which catalyzes the reaction directly upstream of *Bx1* (Figure S10). Two additional genes encoding indole-3-glycerolphosphate synthases are present in *Z. mays (GRMZM2G169516*and *GRMZM2G145870),* but neither was strongly co-expressed with those in the benzoxazinoid pathway. Similarly, the two additional paralogs to *Bx1* in *Z. mays (TSA* and *IGL,* responsible for the production of tryptophan and volatile indole, respectively) formed independent co-expression modules, consistent with their distinct metabolic and ecological roles (Figure S10)^70,71^. The inclusion of an unlinked IGPS gene in the benzoxazinoid co-expression modules suggests that the first committed step in the biosynthesis pathway may start one reaction earlier than previously predicted based on the DIMBOA BGC gene content.

To test whether *GT* and *IGPS* are likely to be involved in benzoxazinoid biosynthesis, we measured their gene expression responses to two different types of insect herbivory (aphid and caterpillar), ecological conditions under which benzoxazinoid biosynthesis genes are typically induced^72,73^. *GT* showed gene expression responses similar to *Bx8* and *Bx9,* being induced within the first few hours after the introduction of insect herbivores (Figure S11). Although the median fold change of expression relative to controls is small (< 5) for all three glucosyltransferases, this result is consistent with a putative role of *GT* in creating stable benzoxazinoid glucosides along with *Bx8* and *Bx9.IGPS* was also significantly induced in response to insect herbivory, mostly notably in the caterpillar feeding experiment in which *IGPS* expression increased over 50-fold during a 24-hour period (Figure S11). In contrast, the two other indole-3-glycerolphosphate synthase genes showed little to no response to herbivory, consistent with this *IGPS* encoding a specialized enzyme involved in benzoxazinoid biosynthesis or volatile indole, which is also induced by caterpillar herbivory^74^.

### Bioinformatically predicted BGCs in plants do not form co-expression modules and are typically not co-expressed

To examine whether putative BGCs (i.e., predicted based on physical clustering and with no known associated products) show evidence of co-regulation in response to specific environmental conditions, we investigated whether they were also recovered in our co-expression network analysis. We found that two different sets of putative BGCs showed little to no co-expression (Figure S12). Specifically, both the 137 Enzyme Commission (EC)-based BGCs predicted by Chae et al.^8^ and the 51 BGCs predicted by the antibiotics and secondary metabolism analysis shell (antiSMASH)^34^ had median MR scores of 9,670 and 10,890, respectively. Furthermore, the EC-based BGCs’ distribution of co-expression was similar to that of the control distribution of neighboring genes (P = 0.187, Wilcoxon rank sum test), whereas the co-expression of antiSMASH BGCs was significantly lower than that of the control (P=0. 027) (Figure 5a and Table S10). In contrast, the six validated BGCs had a median MR score of 17.4 and were significantly more co-expressed than the control (P = 3.20 × 10^−4^) (Figure 5a and Table S10). Similarly, the 13 terpene synthase-cytochrome P450 (TS-CYP) pairs identified by Boutanaev et al.^44^ were variably co-expressed with a median MR score of 45. Although two of the 13 TS-CYP pairs were negatively correlated in their expression, the TS-CYP distribution was still significantly better than the control (P = 2.73 × 10^−4^) (Figure 5a and Table S10).

**Figure 5.**
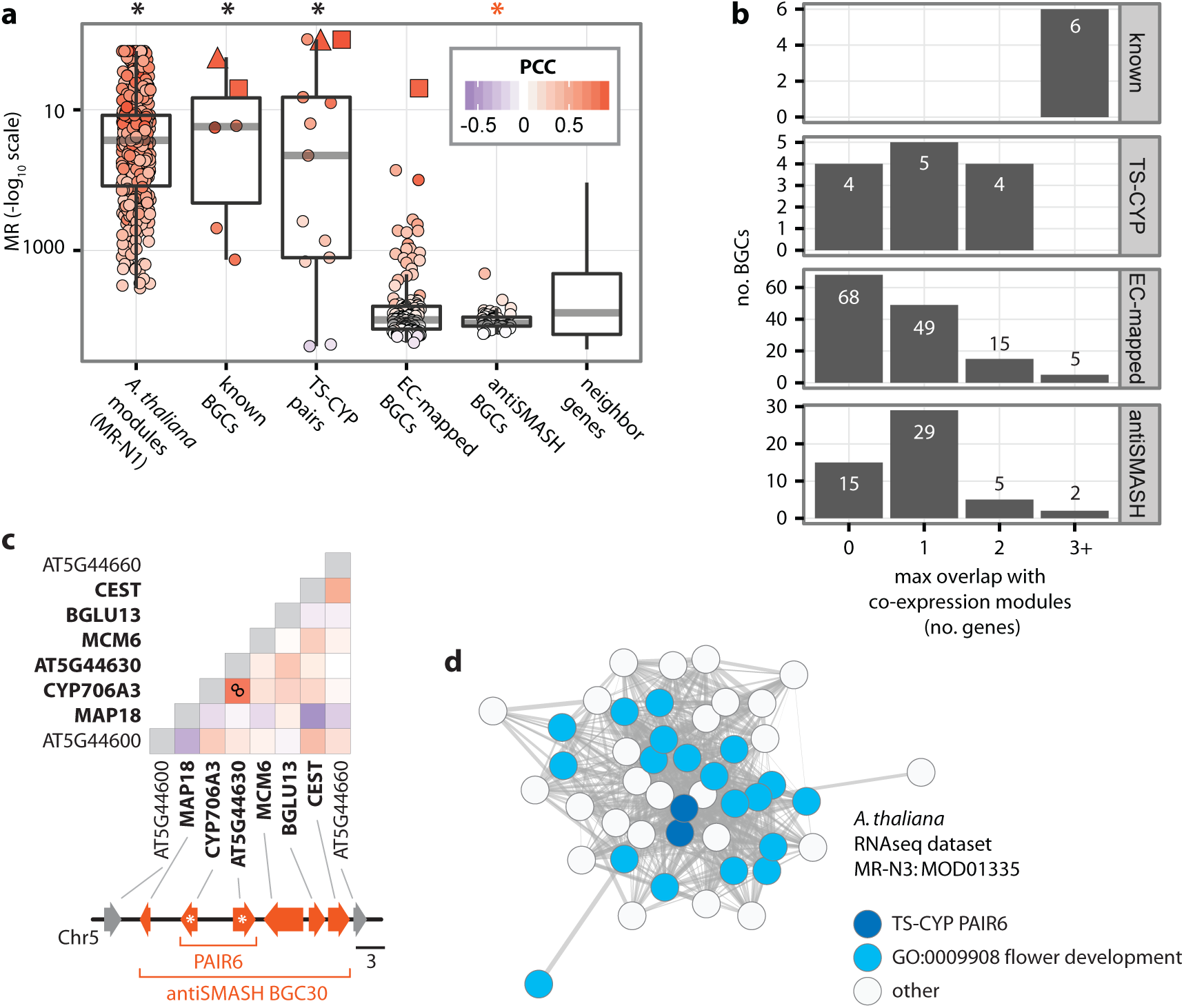
The genes comprising the majority of bioinformatically predicted BGCs are not co-expressed. a) Comparison of average co-expression of modules versus characterized and putative BGCs. The bottom and top of each box plot correspond to the first and third quartiles (the 25th and 75th percentiles), respectively. The lower whisker extends from the box bottom to the lowest value within 1.5 * IQR (Inter-Quartile Range, defined as the distance between the first and third quartiles) of the first quartile. The upper whisker extends from the box top to the highest value that is within 1.5 * IQR of the third quartile. Red squares and triangles indicate BGCs or gene pairs that correspond to the all or part of the thalianol and marneral BGCs, respectively. Asterisks denote significant deviation from the control distribution of neighboring genes; *\*P* ≤ 0.05 (Wilcoxon rank sum tests). b) From top to bottom, histogram of maximum overlap between co-expression modules and known (characterized) BGCs, TS-CYP gene pairs, EC-mapped BGCs, and antiSMASH BGCs. c) Heatmap depicting the correlation of co-expression for a eight-gene region of chromosome five in *A. thaliana* containing an example antiSMASH BGC (BGC30) and TS-CYP gene pair (PAIR6). Diagonal number indicates MR score; squares are blank if MR ≥ 100. Heatmap scale is the same as in part a. BGC genes are bolded in the heatmap and colored red in the accompanying chromosome segments (TS-CYP pair is marked with asterisks). Scale bars are in kilobase pairs. d) Network map of a module that maximally overlaps with BGC30. Overlapping genes (TS-CYP PAIR6) are colored dark blue.

Not surprisingly, given the lack of co-expression, putative BGCs, by and large, did not form co-expression modules, with only 7 / 188 putative BGCs overlapping by three genes or more with co-expression modules. In contrast, 78 / 188 putative BGCs overlapped with co-expression modules by only one gene, indicating that the genes within these BGCs were more strongly co-expressed with genes outside their cluster than with those inside (Figure 5b and Table S8).

An example of the poor association between co-expression modules and putative BGCs comes from the antiSMASH-predicted BGC30. Only 2 / 6 genes in BGC30 showed strong pairwise co-expression: a TS-CYP pair also identified by Boutanaev et al. and labeled PAIR6 (Figure 5c). The terpene synthase (AT5G44630) of PAIR6 is known to be involved in the production of sesquiterpenoid flower volatiles^75^. This functional annotation is supported by our module analysis, which assigned PAIR6 to a co-expression module consisting of 46 physically unlinked genes that are significantly enriched for gene ontology terms associated with flower development (Figure 5d and Table S11). A second example comes from the EC-mapped BGC130. None of the genes in this BGC were strongly co-expressed with each other (Figure S12). Instead, one gene in the BGC, *GSTU20*, is a known participant in metGSL biosynthesis, an association that is recovered by co-expression modules in our analysis (Figure 2 and Table S8).

## Discussion

An enormous number of novel plant SMs awaits discovery and characterization^4^. Yet, due to their rapid evolution and narrow taxonomic distribution^7–9^, SM pathways and genes are often unknown, slowing the pace of discovery. Gene co-expression and chromosomal proximity are two omics-level traits that can be harnessed for high-throughput prediction of SM pathways and genes^4^, but their general utility remained unknown. By examining 10 global co-expression datasets—each a meta-analysis of 172 to 15,275 transcriptome experiments—across eight plant model organisms, we found that gene co-expression was powerful in identifying known SM pathways, irrespective of the location of their genes in the genome, as well as in predicting novel SM gene associations. Below, we discuss why gene proximity may not be a reliable method of SM pathway identification in plant genomes as well as enumerate the advantages and caveats of our co-expression network-based approach.

It is well established that genes in SM pathways are spatially and temporally regulated in response to diverse ecological conditions; arguably, this shared regulatory program is one of the defining characteristics uniting genes belonging to SM pathways^1,11,12^. Furthermore, numerous gene expression studies of the genes participating in diverse SM pathways, including BGCs, from diverse organisms show that SM pathway genes typically share similar gene expression patterns (i.e., they are co-expressed)^21,32,33,64,65,76,77^. Simply put, gene co-expression can be predictive of membership in a given SM pathway. The question then is whether one can employ genome-wide or global gene expression data to predict SM pathways in a high-throughput fashion. The results of our analyses suggest that this is the case; modules in global co-expression networks constructed from genome-wide expression studies across myriads of different conditions in *A. thaliana* were significantly enriched in genes associated with diverse SM-related metabolic functions (Figure 1a). Moreover, modules recovered many experimentally validated SM pathways in these plants (Table S5 and Table S8), including the six known to form BGCs (Figures 4).

It is also well established that gene arrangement in plant genomes is not random^78^. For example, as much as 60% of metabolic pathways in *A. thaliana* (as measured by KEGG) show statistically significant higher levels of physical proximity in the genome than expected by chance^79^. The most extreme version of this “closer than expected” gene arrangement is the growing list BGCs involved in plant SM biosynthesis^42^. While the statistical significance of this pattern is non-debatable, the degree to which gene arrangement is predictive of genes’ participation in the same pathway is not immediately obvious. For example, the genes of many known plant SM pathways^52,80^ do not form BGCs, while other pathways consist of a combination of clustered and unclustered genes^64,69,81^. Complicating matters further, SM pathways may form a BGC in some species but not others^82^. Given that the majority of known plant SM pathways does not form BGCs, it is perhaps not surprising that nearly all putative plant SM BGCs, which were predicted based solely on gene proximity, were not co-expressed (Figure 5).

We interpret this absence of co-expression as evidence that most of these putative BGCs likely do not correspond with actual SM pathways and that gene proximity is insufficient to be used as the primary input for predicting SM pathways in plant genomes. Admittedly, the strength of this argument rests on whether the global co-expression networks that we have constructed accurately capture the spatial and temporal regulation of BGCs in response to the diverse ecological conditions plants experience, which is at least partially dependent on the number and types of the conditions sampled^83^. For example, genes in a BGC or pathway that are never expressed or are not variably expressed across conditions would not be correlated with each other in our analysis. Although this is a valid concern, the hundreds to thousands of conditions^25^ used to construct each co-expression dataset (Table S1) as well as the recovery of many known SM pathways from these organisms (Table S5 and Table S8), suggest that its effect is unlikely to influence our major findings. Going forward, increased resolution of BGCs and SM pathways in co-expression networks will require the inclusion of data from more tissues, time points, and environmental conditions during which SM genes and pathways are likely to vary in their regulation, for example different types of insect herbivory^69,72,84^ and complex field conditions^85^.

Another caveat associated with predicting SM pathways from global co-expression networks is that SM pathways whose expression profiles are highly similar would be predicted to comprise a single pathway. This will likely be a more common occurrence, and examples of this behavior are present in our results. Specifically, the two triterpenoid BGCs in *A. thaliana* were almost always combined in the same co-expression module, regardless of the network investigated (Figure S7); the same was true for the two diterpenoid BGCs in *O. sativa* (Figure S8). Although predicting individual SM pathways is obviously ideal, the lumping of multiple pathways into one may in some cases reveal novel biology. For example, such a pattern could also be indicative of crosstalk between SM pathways or BGCs, or that multiple SM pathways are employed in response to the same set of environmental conditions.

The final caveat is that our approach will not be as powerful in cases where some of the genes in the pathway are not under the same regulatory program as the others. For example, we noted that the genes encoding terminal modification enzymes, such as the genes for side-chain modification of glucosinolates (Figure S4) or the UDP-glucosyltransferases in *S. lycopersicum (GAME2)* and *Z. mays (Bx8-Bx14),* had expression profiles that were quite different from those of core pathway genes; thus, they were often not recovered in the same modules as their corresponding core SM pathway genes. It is possible that additional sampling of appropriate expression conditions could allow for recovery of these terminal metabolic branches in co-expression modules that include the rest of the pathway. However, the terminal SM genes and products can be under balancing or diversifying selection^56^; moreover, the core and terminal steps in an SM pathway may take place in different tissues^86^. In cases like these, the terminal metabolic branches and core SM pathway may be identified as distinct co-expression modules in global co-expression networks no matter how many conditions are sampled.

In summary, our results indicate that generating and constructing global gene co-expression networks is a powerful and promising approach to the challenge of high-throughput prediction and study of plant SM pathways. Global gene co-expression networks can straightforwardly be constructed for any plant, model or non-model, as long as the organism’s transcriptome can be sampled under a range of conditions. In principle, this would not require a genome sequence, only a high quality *de novo* transcriptome assembly. Furthermore, global gene co-expression networks could be used in conjunction with other high-throughput data types (e.g., proteomics, metabolomics). We believe that combining high throughput transcriptomics across ecological conditions with network biology will transform our understanding of the genetic basis and architecture of plant natural products and usher in a new era of exploration of their chemodiversity.

## Materials and Methods

Genome annotations, protein sequences, and gene co-expression values, measured using Pearson’s correlation coefficient (PCC) and mutual rank (MR), across the eight plant species were downloaded from the ATTED-II^25^, ALCOdb^46^, NCBI RefSeq and JGI Genome Portal databases (Table S1). ATTED-II co-expression datasets with less than 50% coverage of the target genome were excluded. The MR score for two example genes A and B is given by the formula,

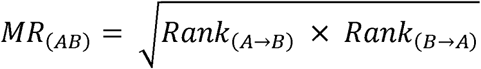

where *Rank*_(*A*→*B*)_ is the rank of gene B in a PCC-ordered list of gene A against all other genes in the microarray or RNAseq meta-analysis; similarly, *Rank*_(*B*→*A*_) is the rank of gene A in a PCC-ordered list of gene B against all other genes, with smaller MR scores indicating stronger co-expression between gene pairs^47^. MR scores were converted to network edge weights using 5 different rates of exponential decay (Figure S1). Any edge with PCC < 0.3 or edge weight < 0.01 was excluded.

Comparison of MR-and PCC-based networks, showed that the MR-based networks were more comparable between species and datasets. For example, PCC-based networks were more sensitive (variable) to differences in the number of experimental samples and genome coverage between datasets in the two species that had microarray- and RNAseq-based datasets *(A. thaliana* and *O. sativa).* In contrast, the MR-based networks were more robust to dataset differences (Figure S13), in agreement with the original description of the MR metric by Obayashi and Kinoshita^47^. Moreover, MR-based networks were remarkably consistent with respect to the number of genes they contained; in contrast, PCC-based networks sometimes varied by orders of magnitude in the number of genes included (Figure S13), Finally, MR-based networks consistently included nearly all genes in a given dataset, regardless of the MR threshold stringency employed; that was not the case with PCC-based networks (Figure S13 and Table S2). For these reasons, we chose to focus the investigation on the MR-based networks.

Modules of tightly co-expressed genes were detected using ClusterONE using default parameters^48^. Modules with ClusterONE *P* value > 0.1 were excluded. Modules were considered ‘SM-like’ if they contained 2 or more non-homologous genes with a significant match to a curated list of PFAM domains present in experimentally verified (Evidence = ‘EV-EXP’) genes assigned to MetaCyc^51^ SECONDARY BIOSYNTHESIS pathways (hmmsearch^87^ using default inclusion thresholds; Table S6). SM-like modules were then binned into meta-modules of non-overlapping gene sets.

Bioinformatically-predicted BGCs were obtained from the published literature^8,44^ and by running the *A. thaliana* reference genome (TAIR10; each protein-coding gene was represented by its longest transcript) through antiSMASH v3.0.4^34^ with the ––clusterblast ––subclusterblast ––smcogs options enabled. Average co–expression of each gene set (module or BGC) was calculated as the average MR score across all gene pairs in the set.

All statistical analyses were performed in R, including dhyper (hypergeometric), wilcox.test (Wilcoxon Rank Sum), p.adjust (Benjamini and Hochberg adjusted *P*-value) from the stats package. Network maps were drawn using a Fruchterman-Reingold force-directed layout using the igraph R package (http://igraph.org).

## Data Availability

All co-expression modules identified in our analysis are included in the supplemental files online (Dataset S1).

## Acknowledgements

We thank members of the Rokas lab and the National Science Foundation’s Plant Genome Research Program for helpful discussions. This work was conducted in part using the resources of the Advanced Computing Center for Research and Education at Vanderbilt University. This material is based upon work supported by the National Science Foundation (http://www.nsf.gov) under Grants IOS-1401682 to JHW, DEB-1442113 to AR, and IOS-1339237 to GJ.

### Author Contributions

Conceived and designed the experiments: JHW VT GJ DJK AR. Performed the experiments: JHW ATB VT. Analyzed the data: JHW ATB VT GJ DJK AR. Contributed reagents/materials/analysis tools: JHW VT GJ AR. Wrote the paper: JHW AR. All authors read, commented on, and approved the manuscript.

## Supplemental files

Dataset S1. Co-expression modules (txt).

Figure S1. Co-expression network pipeline (pdf).

Figure S2. MetaCyc pathway enrichment analysis of experimentally characterized genes in *A. thaliana* (pdf).

Figure S3. Overlapping co-expressed modules recover the pathway for metGSL biosynthesis in *A. thaliana* (pdf).

Figure S4. Comparison of degree of gene co-expression in core versus terminal modification genes in metGSL biosynthesis (pdf).

Figure S5. Maximum likelihood phylogeny of Brassicaceae MAM and IPMS sequences (pdf).

Figure S6. Maximum likelihood phylogeny of Brassicaceae GSTU sequences (pdf).

Figure S7. Network maps of co-expression modules involved in thalianol and marneral triterpenoid biosynthesis in *A. thaliana* (pdf).

Figure S8. Network map of co-expression module involved in momilactone and phytocassane diterpenoid biosynthesis in *O. sativa* (pdf).

Figure S9. Network maps of co-expression modules involved in tomatine biosynthesis in *S. lycopersicum.* (pdf).

Figure S10. Pathway diagram and network map of DIMBOA biosynthesis and related pathways in *Z. mays* (pdf).

Figure S11. Gene expression response to insect feeding in *Z. mays* (pdf).

Figure S12. Coexpression pattern of seven putative BGCs in plants (pdf).

Figure S13. Comparison of Mutual Rank-based and Pearson’s correlation-based networks (pdf).

Table S1. Downloaded datasets (xlsx).

Table S2. Descriptive statistics for co-expression networks (xlsx).

Table S3. *A. thaliana* genes assigned to MetaCyc pathways and pathway ontologies (xlsx).

Table S4. Test for enrichment/depletion of MetaCyc pathway categories and classes in module genes (xlsx).

Table S5. Recovery of MetaCyc pathways in co-expression modules (xlsx).

Table S6. List of Pfam domains found in SM pathways in MetaCyc (xlsx).

Table S7. metGSL biosynthesis genes in *A. thaliana* and *B. rapa* (xlsx).

Table S8. Recovery of metGSL pathways, characterized BGCs, and putative BGCs in co-expression modules (xlsx).

Table S9. List of functionally characterized BGCs in plants with co-expression data on ATTED-II (xlsx).

Table S10. Average co-expression of gene modules, characterized BGCs, and putative BGCs (xlsx).

Table S11. GO enrichment test of a 46-gene *A. thaliana* module involved in flower development (xlsx).

